# Teaching transposon classification as a means to crowd source the curation of repeat annotation – a tardigrade perspective

**DOI:** 10.1101/2023.11.06.565293

**Authors:** Valentina Peona, Jacopo Martelossi, Dareen Almojil, Julia Bocharkina, Ioana Brännström, Max Brown, Alice Cang, Tomas Carrasco Valenzuela, Jon DeVries, Meredith Doellman, Daniel Elsner, Pamela Espindola Hernandez, Guillermo Friis Montoya, Bence Gaspar, Danijela Zagorski, Paweł Hałakuc, Beti Ivanovska, Christopher Laumer, Robert Lehmann, Ljudevit Luka Boštjančić, Rahia Mashoodh, Sofia Mazzoleni, Alice Mouton, Maria Nilsson Janke, Yifan Pei, Giacomo Potente, Panagiotis Provataris, José Ramón Pardos, Ravindra Raut, Tomasa Sbaffi, Florian Schwarz, Jessica Stapley, Lewis Stevens, Nusrat Sultana, Radka Symonova, Mohadeseh Tahami, Alice Urzì, Heidi Yang, Abdullah Yusuf, Carlo Pecoraro, Alexander Suh

## Abstract

The advancement of sequencing technologies results in the rapid release of hundreds of new genome assemblies a year providing unprecedented resources for the study of genome evolution. Within this context, the significance of in-depth analyses of repetitive elements, transposable elements (TEs) in particular, is increasingly recognized in understanding genome evolution. Despite the plethora of available bioinformatic tools for identifying and annotating TEs, the phylogenetic distance of the target species from a curated and classified database of repetitive element sequences constrains any automated annotation effort. Manual curation of raw repeat libraries is deemed essential due to the frequent incompleteness of automatically generated consensus sequences. However, manual curation and classification are time-consuming processes that offer limited short-term academic rewards and are typically confined to a few research groups where methods are taught through hands-on experience. Crowd sourcing efforts could offer a significant opportunity to bridge the gap between learning the methods of curation effectively and empowering the scientific community with high-quality, reusable repeat libraries. Here, we present an example of such crowd sourcing effort developed through both in-person and online courses built around a collaborative peer-reviewed teaching process that can be used as teaching reference guide for similar projects. The collaborative manual curation of TEs from two tardigrade species, for which there were no TE libraries available, resulted in the successful characterization of hundreds of new and diverse TEs: A hidden treasure awaits discovery within non-model organisms.

## Background

The importance of in-depth analyses of repetitive elements, particularly transposable elements (TEs), is becoming more and more fundamental to understand genome evolution and the genetic basis of adaptation [1]. While there is a wealth of bioinformatic tools available for the identification and annotation of TEs (https://tehub.org/en/resources/repeat_tools), any automated annotation effort is limited by the phylogenetic distance of the target species to a database of curated and classified repetitive element sequences [2]. For example, in birds where zebra finch and chicken have well-characterized repetitive elements because their genomes were first sequenced in large consortia during the pre-genomics era [3,4], automated annotation of other bird genomes will render most repeats as correctly classified [5,6]. On the other hand, in taxa as diverse and divergent as insects, up to 85% of repetitive sequences can remain of “unknown” classification in non-*Drosophila* species [7]. This is problematic. Inferences about the mobility and accumulation of TEs, as well as their potential effects on the host, are not feasible for unclassified repeats, as well as for incorrectly classified repeats if the automated classification is based on short, spurious nucleotide sequence similarity [8,9].

The reference bias in TE classification reflects the history of the TE field in the genomics era: In the 1990s and 2000s, there were usually multiple people tasked with TE identification, classification, and annotation for each genome project, yielding manually curated consensus sequences (namely representative sequences which quality was controlled and improved) and fully classified TE libraries deposited in databases such as Repbase [2]. Over the last ten years, however, the number of genome projects both of individual labs as well as large consortia has increased exponentially and so have speed and number of automated TE annotation efforts [10–12], while time and personnel have remained limited for curated TE annotation efforts. Similar to taxonomic expertise required for identifying and classifying organisms, TE identification and classification need hands-on experience with manual curation for months or even years per genome [1] which is usually taught through knowledge passed within genome projects and research groups. Recent efforts [13–15] have started to make manual curation accessible to a broader scientific audience, with the aim to increase reproducibility and comparability. However, what cannot be changed is that there are hundreds if not thousands of genomes per TE-interested researcher with more or less pressing priority for time-consuming manual curation.

Low scalability and people power are major obstacles that need to be overcome by the many facets of computational biology where curation is essential. Annotation efforts of other genomic features have shown that crowd sourcing through teaching [16–22], or “course sourcing” as we call it, has the benefit of providing participants with hands-on skills for curation and experience on how to reconcile biology with technical limitations, while simultaneously sharing the workload of time-consuming curation across multiple people working on different parts at the same time. Thus, we argue that a TE curation effort that would take months or years for a single person may fit into a few days or weeks of teaching, of course as long as reproducibility and comparability are ensured throughout course duration.

Here, we present our “course sourcing” experience from two iterations of a Physalia Course on TE identification, classification, and annotation. We focused on two species of tardigrades as a case study to motivate student-centered learning through direct contribution to scientific knowledge: Tardigrades are, to our knowledge, the most high-ranking animal phylum without curated TE annotation, very clearly illustrated by the fact that in previous genome analyses, almost all repeats remained of “unknown” classification [23]. Tardigrades are a diverse group of aquatic and terrestrial animals which show extraordinary ability to survive extreme environments by entering the state of cryptobiosis [24]. This animal clade comprises almost 1,200 described species belonging to Panarthropoda [25] and the two species used in the courses are closely related and belong to the Hypsibiidae family [23].

The first course took place in person in June 2018 in Berlin across five full-time work days: The first three days familiarized the 13 participants with the biology of TEs, concepts for classification, and methods for annotation using the tardigrade *Hypsibius dujardini*, while the last two days had a student-centered learning format where each participant was able to deepen knowledge where needed and curate as many TEs as possible from the target species. The second course took place virtually in June 2021 due to the Covid-19 pandemic and comprised five afternoons in the Berlin time zone to minimize Zoom fatigue. The overall format was similar to the prior in-person course but with 24 participants and focusing on another tardigrade, *Ramazottius varieornatus*, which the participants identified to have not a single shared TE family with the tardigrade *H. dujardini* curated in the 2018 course. Between the two courses, the participants were able to uncover a vast diversity of TEs and successfully curate almost 500 consensus sequences. We demonstrate therefore that a collaborative approach is a valuable means to achieve significant results for the scientific community and we hope to share with the community a teaching reference for future similar efforts, because: A hidden treasure always awaits discovery in non-model organisms.

## Results and Discussion

Incorporating crowd sourcing efforts within a classroom setting (“course sourcing”) can represent an invaluable opportunity for teaching, while simultaneously contributing to the scientific community. However, course sourcing also presents its own unique challenges, particularly in terms of minimizing errors, maximizing reproducibility and student engagement. Drawing from our experience in both in-person and virtual settings, we identified several crucial factors in teaching TE manual curation that must be considered during the organization and supervision of such course, like: a) establishing a standardized approach for curation and classification of TE consensus sequences; b) implementing a peer-review process between participants to check on the quality of the curated consensus sequence; c) maintaining meticulous version control of the libraries. Here, we describe how we addressed these points. First, to establish a standard approach to manual curation, we implemented methods widely used in the TE community that have been recently reviewed in detail [13,14]. The approach, briefly, consists in producing and inspecting multi-sequence alignments for each of the consensus sequences automatically generated by RepeatModeler [10]. Each nucleotide position of the “alignable part” of the alignment is carefully inspected to identify the correct termini of the TE while correcting for any ambiguous base or gap. To correct for ambiguous bases, we applied a majority rule and assigned the most representative IUPAC nucleotide character for each position in the alignment (see **Methods**). To correct the consensus sequences where gaps of different lengths are present, we considered each insertion/deletion length as independent events so that a majority rule was applicable to these regions as well. When very complex regions could not be unambiguously solved, stretches of 10 N nucleotides were inserted as placeholder (gap) in the consensus sequence. The TE classification followed the nomenclature used by RepeatMasker to ensure direct compatibility with the tool and its suite of scripts for downstream analysis. Second, when participants completed the curation of their consensus sequences, then their results would go through a peer-review process where both the quality of the sequence and its classification were revised by other participants (or course faculty). During the in-person edition, a random set of consensus sequences curated by one participant was assigned to another participant, while in the second online edition, all sequences were reviewed by the two instructors and one participant (**Figure 1**). The review of the TE sequences continued after the official conclusion of the course. To ensure reproducibility and the documentation of the entire decision-making process for classification, all steps and details of classification were recorded in a shared Google Sheet. The tables would include the changes in consensus sequence names, names of the curators and reviewers and additional comment (**Figure 1**, **Table S1**). Whenever a change was introduced in a consensus sequence (either in the nucleotide sequence itself or in the classification), the new version was directly added to the multi-sequence alignment file used for curation together with the original one. Keeping all the versions of a consensus in the same alignment file and respective notes in the tables allows the implementation of a basic version control useful to check on the steps leading to a particular decision. From the reiteration of the course, we noticed three particularly challenging points for beginners that need an extra supervision effort. The most challenging points are the identification of the correct termini, target site duplications (a hallmark of transposition for the vast majority of TEs) if any, and the correct spelling of the TE categories for classification in accordance with the RepeatMasker nomenclature rules. The last point is of particular importance especially if the repeat annotation is visualized as a landscape using the RepeatMasker scripts (e.g., calcDivergence.pl and createRepeatLanscape.pl) to not cause computing errors and downstream misinterpretations.

**Figure 1.**
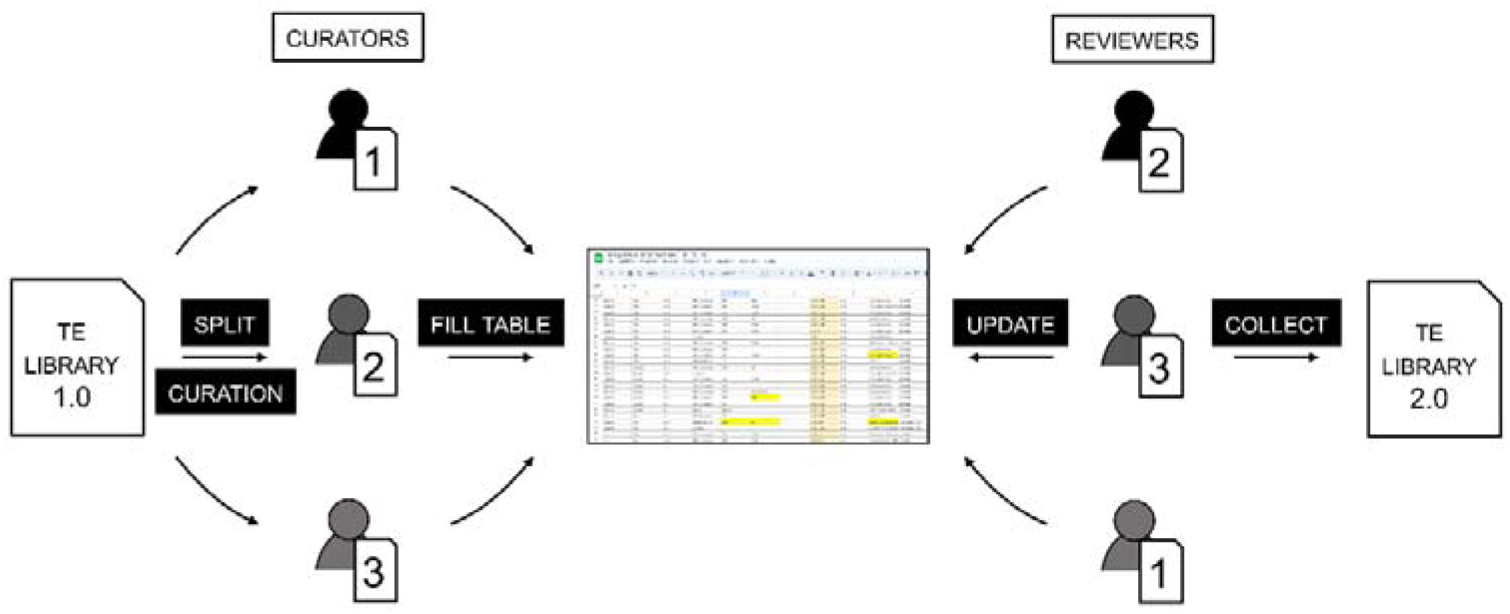
Schematic representation of the peer-reviewed process of TE curation.

Finally, all the tutorials to obtain and curate a TE library are available on the GitHub repository linked to this paper: https://github.com/ValentinaPeona/TardigraTE.

### Improvement of the transposable element libraries

To generate the TE libraries, we first ran RepeatModeler and RepeatModeler2 on both species and obtained 489 and 900 consensus sequences for *H. dujardini* and *R. varieornatus* respectively (**Table 1**). Then the course participants manually curated as many consensus sequences as possible. In about three course days plus voluntary efforts by some participants after each course, the participants were able to curate 286 consensus sequences (58%) of the *H. dujardini* library and 145 consensus sequences (16%) of the *R. varieornatus* library (**Table S1-3**). Given the lack of previously curated libraries from closely related species, most of the consensus sequences were automatically classified as “unknown” by RepeatModeler, but the thorough process of manual curation successfully reclassified 305 unknown consensus sequences (out of a total of 431 curated sequences, 71%) into known categories of elements. After manual curation, we found that most of the two species’ libraries are comprised of DNA transposons and a minority of retrotransposons (**Table 1**). Since many consensus sequences remained uncurated and unclassified, it is possible that the relative percentages of the categories change in the future, but we expect, especially from the composition of the *H. dujardini* library, to mostly find additional (non-autonomous) DNA transposons among the unclassified.

**Table 1:** Overview of classification of tardigrade repeats in the curated libraries. The libraries here described contain both curated and uncurated consensus sequences.

| Species | DNA | LINE | LTR | SINE | Unknown |
| --- | --- | --- | --- | --- | --- |
| *Hypsibius dujardini* | 247 | 12 | 29 | 2 | 199 |
| *Ramazzottius varieornatus* | 203 | 35 | 11 | - | 651 |

The process of manual curation improved the overall level of TE classification of the libraries but also the quality of the individual consensus sequences by correctly identifying their termini and in general by extending their sequence. Indeed, by comparing the lengths of the consensus sequences for the same element, we can notice a marked increase in length after curation (**Figure 2**).

**Figure 2.**
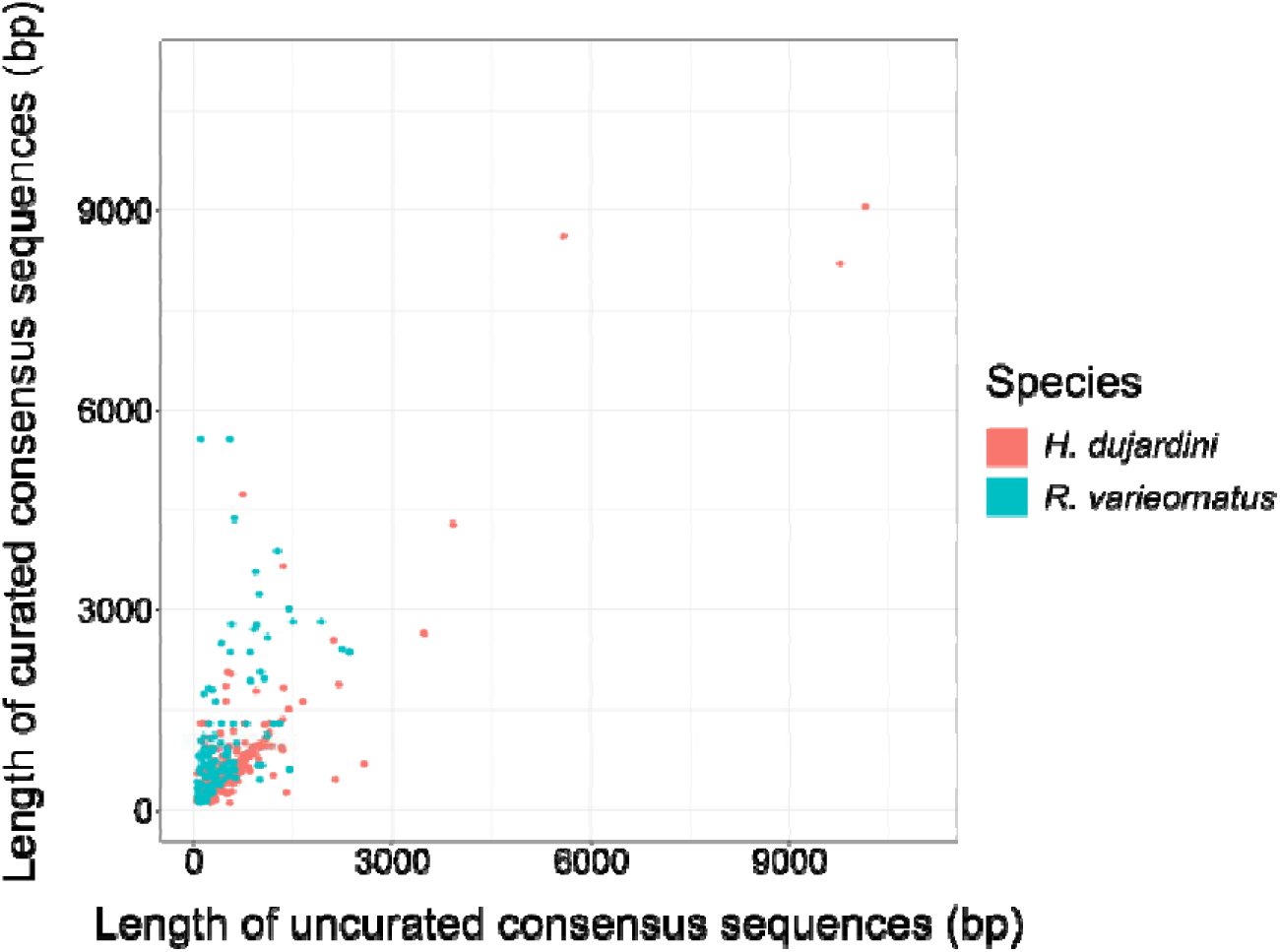
Comparison of the length of the consensus sequences before and after manual curation.

### Diversity of transposable elements

When looking at the diversity of repeats in the partially curated libraries (libraries comprising both curated and uncurated consensus sequences), we identified a total of 419 Class II DNA consensus sequences belonging to the superfamilies/clades CMC, MULE, TcMar, Sola, PiggyBac, Tc4, PIF-Harbinger, Zator, hAT, Maverick, and P. Many of these elements are non-autonomous and show a remarkable diversity of internal structures (**Figure 3**). For Class I retrotransposons, we found 40 LINEs belonging to the superfamilies/clades CR1, CRE, R2, R2-NesL, L2, RTE-X and RTE-BovB and other 35 LTRs belonging to the superfamilies/clades DIRS, Gypsy, Ngaro and Pao.

**Figure 3.**
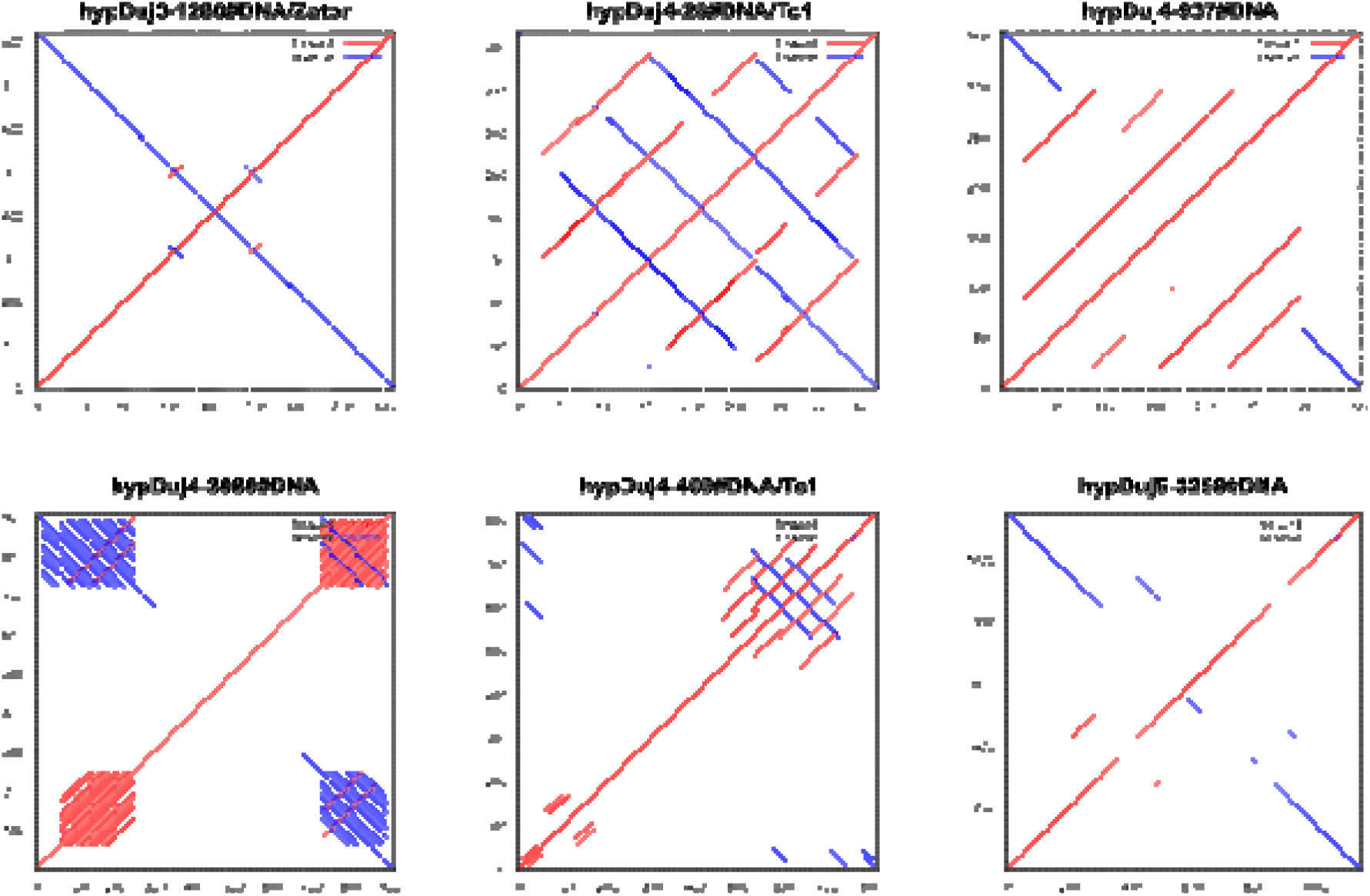
Dotplots of six DNA transposons from the library of *Hypsibius dujardini* produced with the MAFFT online server. These elements were selected by course participants for aesthetic reasons.

To highlight the importance of generating and using custom repeat libraries for the organisms of interest as well as their curation, we masked the two tardigrade genomes and compared how the annotation and accumulation patterns change when using general repeat libraries (in this case the Repbase library for Arthropoda) and species-specific ones before and after curation (**Figure 4**, **Table 2**). The use of the known repeats for Arthropoda available on Repbase provided a poor and insufficient annotation for both species (all the following percentages are given for *H. dujardini* and then for *R. varieornatus*) where only 1.95% and 0.26% of the assemblies were annotated as interspersed repeats and the accumulation patterns were characterized only by likely old insertions. Then the use of species-specific, albeit uncurated, libraries completely changed the percentage of TEs annotated (16.38% and 15.66%) and their accumulation patterns that showed many recently accumulated insertions.

**Figure 4.**
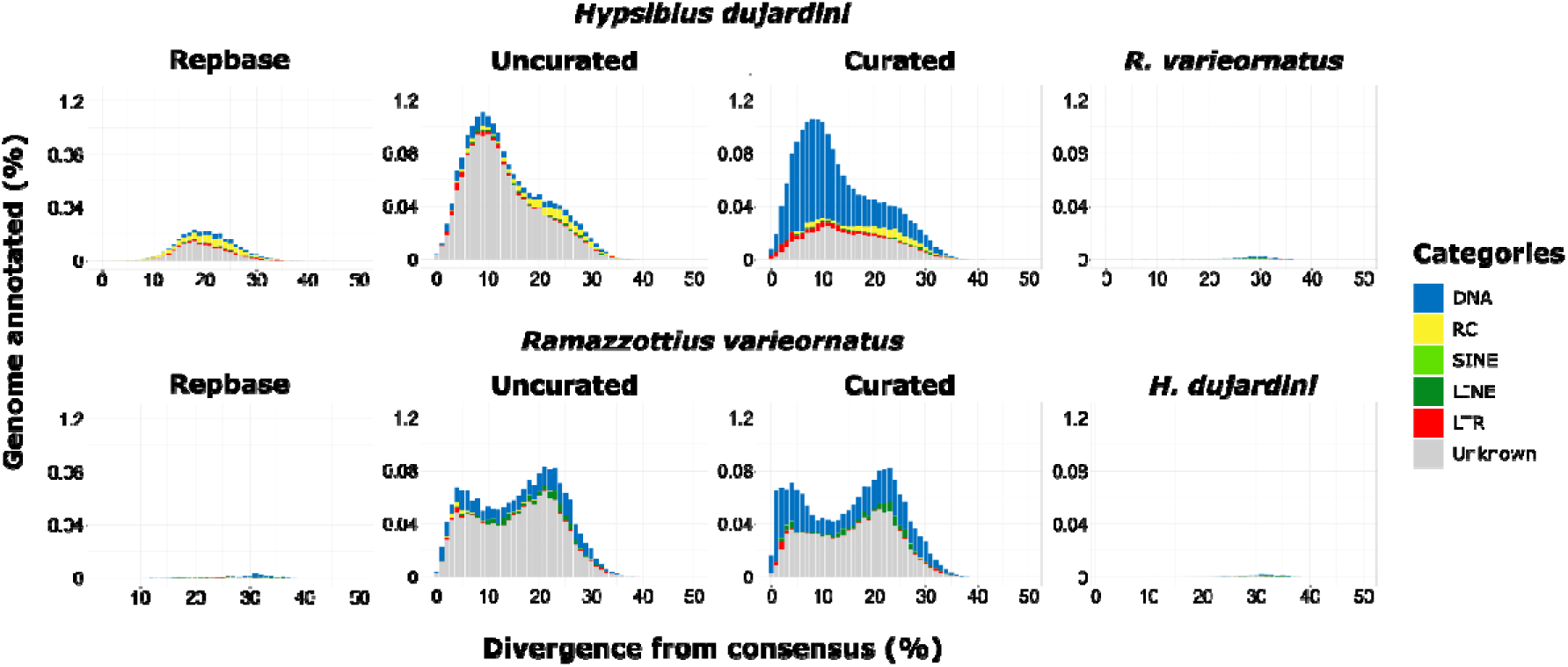
Repeat landscapes of the genomes of *H. dujardini* and *R. varieornatus* annotated with the Repbase (Arthropoda clade), uncurated and curated of both tardigrades combined libraries, and with libraries of the reciprocal species (only species-specific repeats). The divergence from consensus calculated with the Kimura 2-parameter distance model is shown on the x-axis. The percentage of genome annotated is shown on the y-axis.

**Table 2.**
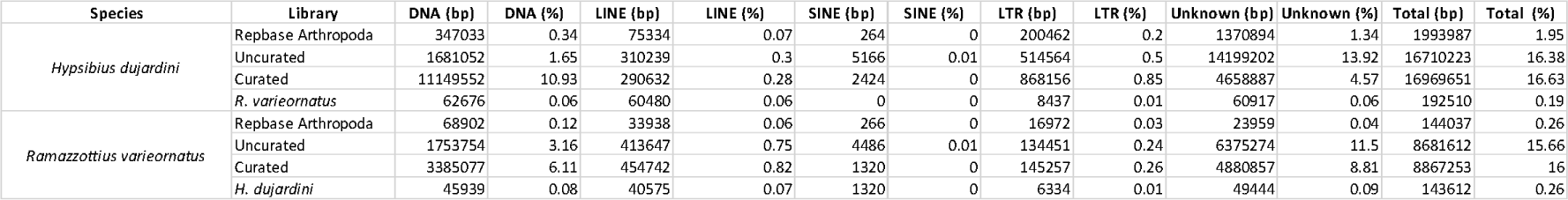
Number of base pairs annotated and percentages of the main TE categories.

While the shape and percentages of the repeat landscapes did not drastically change after the manual curation of the libraries, the curated libraries clearly highlighted a large accumulation of DNA transposons in recent and ancient times alike that were either not present in the other landscapes or were hidden among the “unknown” repeats. Especially for *R. varieornatus*, the curation highlighted a higher accumulation of repeats in the very recent times (1-5% of divergence). This higher accumulation of DNA transposons in recent times is also in line with the finding of multiple putatively active transposable element subfamilies (**Table 3**). Finally, the use of the repeat library of one species to annotate the other species (reciprocal masking) resulted to be almost as insufficient as the use of the Repbase library for Arthropoda stressing once again how important it is to have a capillary knowledge of the repeatome for correct biological interpretations.

**Table 3:** List of repeat subfamilies with putatively ongoing activity, i.e., at least 10 copies with 0% distance to consensus.

| TE category | *Hypsibius dujardini* | *Ramazzottius varieornatus* |
| --- | --- | --- |
| DNA transposon | 7 | 3 |
| LTR retrotransposon | 3 | 0 |
| Unknown | 0 | 2 |

As a demonstrative example of the contribution of the collaborative curation process in providing novel insights into TEs diversity, taxonomic distribution and biology, we decided to deeply characterize consensus sequences that we classified as Tc4. These elements have a rather limited taxonomic distribution, few references in bibliography exist, and they incompletely duplicate the target site upon transposition [26] which can impose challenges for their classification. The Tc4 transposons are DDD elements firstly discovered in *Caenorhabditis elegans* [26] where they recognize the interrupted palindrome CTNAG as target site for insertion, and cause duplication of only the central TNA trinucleotide. Regarding their taxonomic distribution, consensus sequences for Tc4 elements are known and deposited only for nematodes and arthropods in RepeatPeps, Repbase and DFAM. Phylogenetic analyses based on DDD segments confidently placed the four tardigrade Tc4 consensus sequences identified in *R. varieornatus* within the Tc4 clade in a sister relationship with arthropod elements and with a branching pattern that reassemble the Panarthropoda group (tardigrades + onychophorans + arthropods) within Ecdysozoa [27] (**Figure 5A**). The DDD catalytic domain resulted to be highly conserved between different phyla (**Figure 5B**) and the target site of tardigrades mirror what was previously observed in nematodes (i.e., C|TNA|G where “|” marks the transposase cut site **Figure 5C-D**). We could therefore hypothesize that these elements first originated during the diversification of Ecdysozoa. However, broader comparative analyses involving more early-diverging Metazoa clades are necessary to confirm this lineage-specific origin.

**Figure 5.**
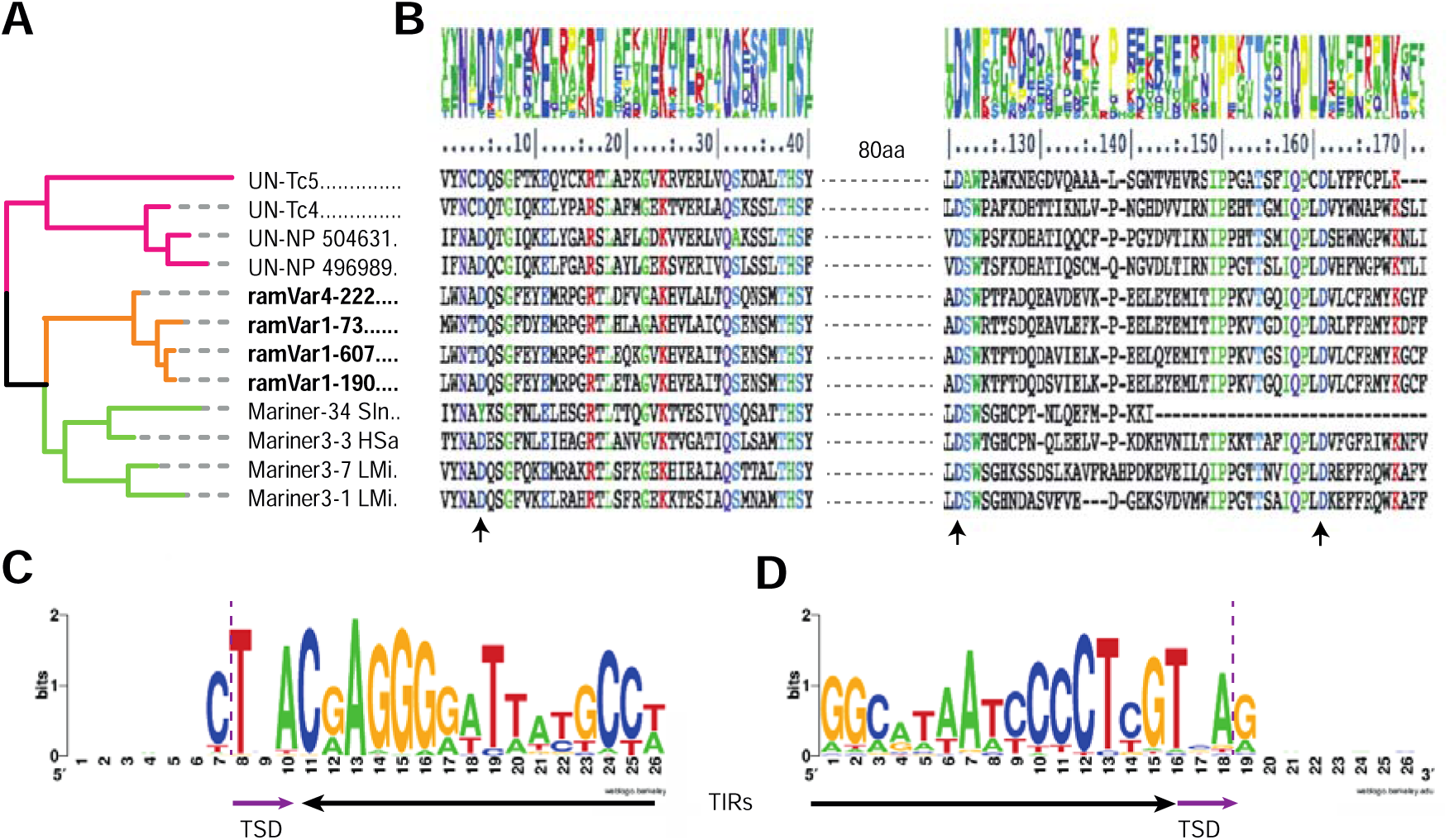
Characterization and phylogenetic analyses of Tc4 elements. (A) Phylogenetic tree of Tc4 consensus sequences based on DDD catalytic domains identified in the *R. varieornatus* consensus sequences, highlighted in bold and orange, together with representative sequences extracted from the RepeatPeps library from nematodes (pink) and insects (green). All nodes received maximal support value. (B) Alignment of DDD catalytic domains of sequences included in phylogenetic analyses. Residues conserved in more than 80% of the sequences are colored. Arrows highlight catalytic DDD residues. Sequence logos of 5’ (C) and 3’ (D) ends of Tc4 elements used to curate the *R. varieornatus* consensus sequences. Black and purple arrows denote terminal inverted repeats (TIRs) and target site duplications (TSDs), respectively. The purple dotted line marks the transposase cut on the CTNAG target site.

### Contributions from the course participants

During both editions of the course, participants were free to explore their favorite topics within the scope of the syllabus and we here share two contributions developed by the participants that can be useful for the entire community. First, an additional repeat library of 130 consensus sequences (119 of which are DNA transposons) was produced with the use of REPET for *R. varieornatus* (**Table S4**). Second, a guide for the classification of TEs from multisequence alignments (**File S1**) that can be a useful starting point for beginners and complementary to more extensive guides [13,14].

## Conclusion

As shown here and in many other studies, repeat annotation is key to correctly identify and interpret patterns of genome evolution and proper annotation is based on a thorough curation of the repeat libraries [8,9,28]. However, it is hard for curation efforts to keep up with the sheer amount of genomes released every year as curation done by single laboratories may require months or even years for a single genome. Recent developments of machine learning-based tools to automatize the curation and classification processes are promising [29–32] and there are additional tools to facilitate the curation process like TEAid [13] and EarlGrey [33]. Until fully automatized, reliable tools are developed and there are manual curation training sets for understudied taxa, we emphasize the need to implement manual curation for repeat libraries as well as to find alternative ways to deal with the curation of hundreds of new libraries. Here we presented one such alternative approach, namely a peer-reviewed course sourcing effort designed to be as reproducible and comparable as possible and where the hands-on tutorials were designed to be meaningful for the participants because they dealt with real unexplored data and directly contributed to the scientific community. The two iterations of this course sourcing effort resulted in the successful curation of hundreds of new and diverse TEs and we hope that this experience and teaching framework can be of use for the genomic and TE communities and to be applicable to other types of data/analysis that need manual curation (e.g., genome assemblies [21,22] and gene annotations).

## Materials and Methods

### Genome assemblies

For this study, we used the genome assemblies of the two tardigrade species: *Hypsibius dujardini* (GCA_002082055.1) and *Ramazzottius varieornatus* (GCA_001949185.1) produced by sequencing a pool of male and female individuals by Yoshida et al. [34]. The *Hypsibius dujardini* genome was assembled using long PacBio and short Illumina reads whereas the *Ramazzottius varieornatus* genome was assembled using a combination of Sanger and Illumina reads [34].

### Raw repetitive element library

To start the *de novo* characterization of transposable elements, we ran RepeatModeler on *H. dujardini* and RepeatModeler2 on *R. varieornatus* [35] using the option -LTR_struct and obtained a library of raw consensus sequences for each of the genomes. RepeatModeler and RepeatModeler2 automatically named the consensus sequences with the prefix “rnd” that we replaced with the abbreviations of the species names: “hypDuj” for *H. dujardini* and “ramVar” for *R. varieornatus*.

The two libraries were then compared to find similar sequences belonging either to the same family or subfamily by using, respectively, the 80-80-80 rule [36] and the 95-80-98 rule [37]. The comparison was done by masking the library of *R. varieornatus* with the library of *H. dujardini* using RepeatMasker [38].

### Manual curation of the consensus sequences

After the generation of the libraries of raw consensus sequences, we proceeded with the collaborative peer-reviewed manual curation step. The participants of the course were split into ten groups and each group received about 80 consensus sequences to curate.

The first step of the curation consisted in the alignment of the raw consensus sequences to the genome of origin using BLAST [39]. The best 20 BLAST hits were selected aligned with their raw consensus sequence with MAFFT [40] which produced a multisequence alignment for each consensus sequence ready to be manually curated (script RepeatModelerPipeline.pl).

Each of the multisequence alignment was then inspected to: 1) find the actual boundaries of the repetitive elements; 2) build a new consensus sequence with Advanced Consensus Maker (https://hcv.lanl.gov/content/sequence/CONSENSUS/AdvConExplain.html); 3) fix ambiguous base and gap calls in the new consensus sequence following a majority rule; 4) find sequence hallmarks to define the repetitive elements as transposable elements (e.g., target site duplication, long terminal repeats, terminal inverted repeats or other motifs). Every new consensus sequence was reported in a common Excel table (**Table S1**). To quantitatively measure the improvement of the repeat libraries after manual curation, we compared the length of consensus sequences before and after curation.

In all the figures and tables, the term “curated” indicates that the library mentioned contains manually curated consensus sequences as well as all the consensus sequences that remained uncurated. Finally, we consider each consensus sequence as a proxy for a transposable element subfamily. However, the consensus sequences were not checked for redundancy and not clustered into families and subfamilies using the 80-80-80 or 95-80-98 rules for nomenclature because the focus of the study was on classifying the consensus sequences into superfamilies and orders of transposable elements.

The code used to produce the consensus sequences and their alignments is provided as tutorial on the GitHub repository https://github.com/ValentinaPeona/TardigraTE.

### Classification

The new consensus sequences were classified using sequence characteristics retrieved by the alignments (e.g., target site duplications, terminal repeats) and homology information retrieved through masking the sequences with Censor [41,42] following the recommendations from [36] and [43]. When the information retrieved by the alignments and Censor were not enough to provide a reliable classification of the elements, the sequences were further analyzed for the presence of informative protein domains using Conserved Domain Database [44–46].

Since the course participants in general had never curated transposable element alignments before, we decided to implement a peer-review process. For the first course (*H. dujardini*), the results of each participant were sent to another participant to check the curated alignments and independently retrieve key information for the classification. The independent sequences and classifications would be compared and fixed if necessary. In the second course (*R. varieornatus*), all sequences were inspected by the same 3 reviewers only who applied the same process as previously described.

### Comparative analysis of the repetitive content

The genome assemblies of both tardigrade species were masked with RepeatMasker 4.1.10 using four different types of TE libraries: 1) known Arthropoda consensus sequences from Repbase; 2) raw uncurated consensus sequences from the respective species; 3) curated consensus sequences together with the consensus sequences that were not curated from the respective species; 4) curated consensus sequences together with the consensus sequences that were curated from the other species. The RepeatMasker output files were then used to get the percentages of the genomes annotated as TEs and to visualize the landscapes of the accumulation of repeats.

Finally, we estimated the number of putative active transposable elements in the two genomes by filtering the RepeatMasker annotation for elements that show at least 10 copies with a 0% divergence from their consensus sequences.

### Characterization of Tc4 elements

During the manual curation process, participants found types of DNA transposons that are currently considered to have a rather restricted phylogenetic distribution like Tc4 Mariner elements, therefore more in-depth analyses were run on these elements. The protein domains of known Tc elements were compared to the Tc4 consensus sequences from the tardigrade species and phylogenetic relationships were established.

Protein homologies of the partially curated repeat libraries were collected using BlastX (e-value 1e-05) [47] against a database of TE-related protein (RepeatPeps library) provided with the RepeatMasker installation. We extracted the amino acid translation of each hit on Tc4 elements based on the coordinates reported in the BlastX output. Resulting protein sequences were aligned together with all members of the TcMar superfamily present in RepeatPeps library using MAFFT (*L-INS-i* mode) [48] and the alignment was manually inspected to identify and isolate the catalytic DDD domain. The resulting trimmed alignment was used for phylogenetic inference with IQ-TREE-2 [49], identifying the best-fit evolutionary model with ModelFinder2 and assessing nodal support with 1000 UltraFastBootstrap replicates [50]. The resulting maximum likelihood tree was mid-point rooted and the Tc4 subtree extracted for visualization purposes. The DDD segments of Tc4 elements were re-aligned using T-Coffee in *expresso* mode [51] to produce conservation scores. A sequence logo of 5’ and 3’ boundaries of identified Tc4 elements was produced extracting all sequences used to curate the four *R. varieornatus* Tc4 elements and keeping the first 15 bp and 11 bp before and after the terminal inverted repeats (TIRs), respectively.

### Additional transposable element library

Participants ran REPET tool V3.0 [52] to produce a de novo transposable element library for *R. varieornatus* in parallel to the one generated by RepeatModeler2. A custom TE library composed by repeats from Repbase and from *H. dujardini* was used to aid REPET in the classification process. Only consensus sequences that showed two or more full-length copies in the *R. varieornatus* genome were retained in the new library. Furthermore, the consensus sequences were scanned for protein domains and presence of TIRs or long terminal repeats (LTRs).

## Abbreviations

LTR: Long Terminal Repeats
TE: transposable element
TIR: Terminal Inverted Repeats

## Declarations

### Ethics approval and consent to participate

Not applicable.

### Consent for publication

Not applicable.

### Availability of data and materials

All data generated or analyzed during this study are included in this published article and its supplementary information files. All newly curated repeat consensus sequences were deposited in Dfam. The code for the tutorials used in the course as well as for the analysis of the manuscript can be found on GitHub: https://github.com/ValentinaPeona/TardigraTE.

### Authors’ contributions

AS conceived the project and VP contributed to its development. VP and JM analyzed the data. AS, VP, JM wrote the manuscript, and all authors revised the manuscript. MT, AM, DA, JS, GP provided additional contributions to the teaching material. All authors except CP contributed to the curation of the repeat library. CP provided and maintained the computational infrastructure during the courses. Authors are listed in alphabetical order.

## Acknowledgements

Part of the analysis were performed on resources provided by the Swedish National Infrastructure for Computing (SNIC) through Uppsala Multidisciplinary Center for Advanced Computational Science (UPPMAX) and CSC-IT Finland.

