## Supplementary material for "Teaching transposon classification as a means to crowd source the curation of repeat annotation – a tardigrade perspective": File_S1

**An identification key for TEs curation**

This guide has been developed by the participants to the 2021 Physalia course and it is meant to be an introduction to the classification process of transposable elements using multi-sequence alignments. Tutorials for how to obtain such alignments are present in the GitHub repository https://github.com/ValentinaPeona/TardigraTE

This guide can be of particular use as companion to more extensive guides about manual curation as Goubert et al 2022 (https://doi.org/10.1186/s13100-021-00259-7) and Storer et al 2021 (https://doi.org/10.1002/cpz1.154).

**Glossary:**

- TE: Transposable Element
- TSD = Target Site Duplication, a short direct repeat that is generated on both flanks of a TE upon insertion.
- LTR = are Long Terminal Repeats that range in size from 100 bp to 25 kb. The LTR flanking region range from 100 bp to 5 kb. Upon integration, LTR produce TSD of 4-6 bp. LTRs typically contain ORFs for GAC, a structural protein for virus-like particles and for POL (POL encodes an aspartic proteinase (AP), reverse transcriptase RNase H (RH), and DDE integrase (INT).
- TIRs = Terminal Inverted Repeats of variable lengths present in DNA transposon class II
- Autonomous TEs: An element that encode all the domains that are typically necessary for its transposition, without implying that the element is either functional or active.
- Autonomous but defective TEs: defective due to mutation(s).
- Non- autonomous TEs: Any group of elements that lacks some or all of the domains found in autonomous elements. Occasionally have highly degenerative coding region or even completely lack coding capacity. Some non- autonomous TEs lack some genes but contain others, for example, Caspar family (Super family CACTA) often lacks the transposase gene but still contains the second ORF. Whereas BARE2 elements in the Triticeae have a conserved deletion that inactivates “gag”. Non-autonomous TEs: classified based on significant similarities of their terminal inverted repeats and target site duplication to those in known autonomous DNA transposons.

**Visual general guide illustrating the main classification of TEs:**

A) **Classification of TEs:**

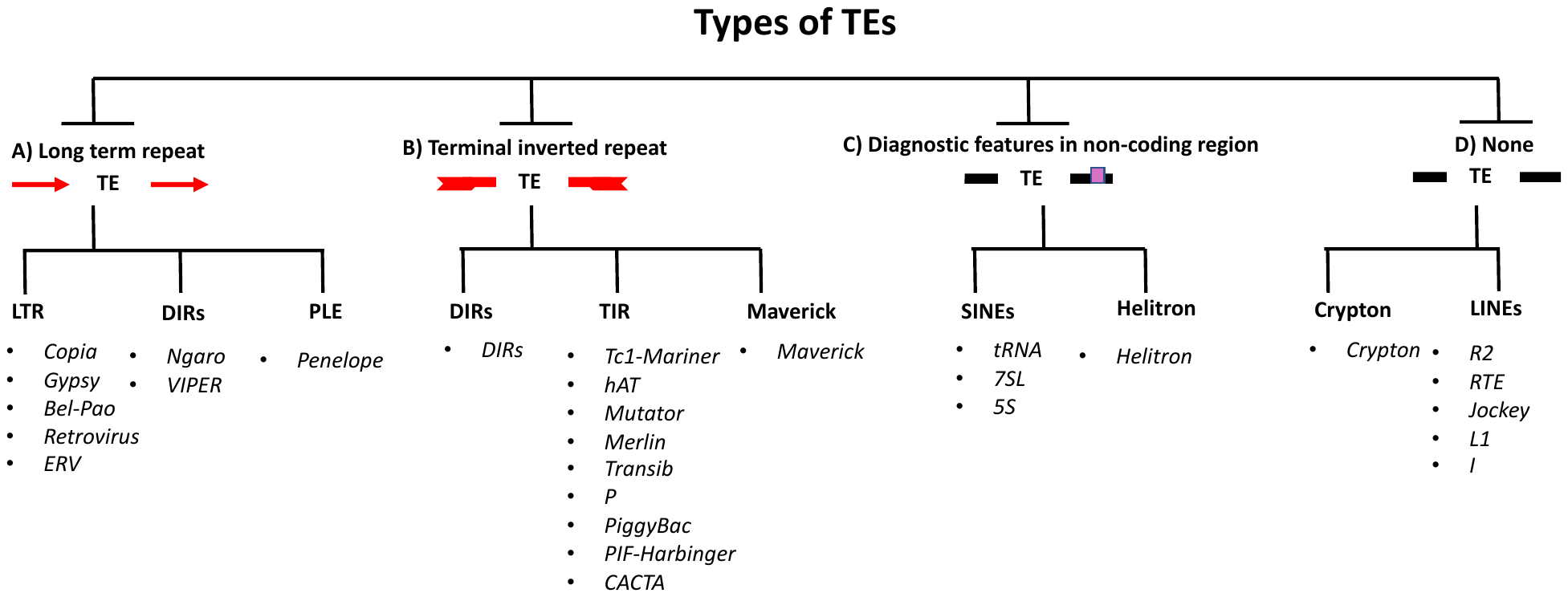

B) **DNA sequence features for each TE class:**

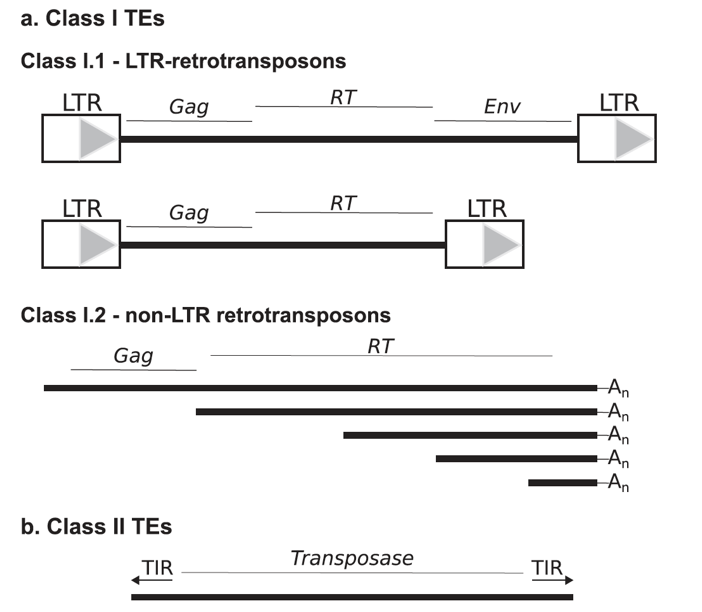

Adapted from Piegu et al (2015), who reproduced it from Finnegan (1992.) showing a proposal for TE classification based on TE DNA sequence features. **(a) Class I** is composed of elements that transpose by reverse transcription of an RNA intermediate and includes two sub-groups. Class I TEs have all the signatures of an endogenous retrovirus-like element, including long terminal repeats at both ends and open reading frames (ORFs) coding for a group antigen (Gag), a reverse transcriptase (RT), and in some case an envelope protein. Class I.2 elements only have Gag and RT ORFs. Class I.2 is composed of elements that look like long retro-inserted messenger RNA (mRNA) with an A-rich tail at their 3’ end. Within a species of such elements many copies are truncated at their 50 ends. **(b) Class II** elements that transpose directly from DNA to DNA and have short terminal inverted repeats (arrowed) at both ends. They contain a gene coding a transposase, an enzyme required for their own transposition.

**Guide to classify transposable elements from multi-sequence alignments**

1) The 5’ and 3’ ends of the alignable regions or consensus sequence have long terminal repeat starting with TG on the 5’ and ending with CA on the 3’ end (e.g., 5’ – TG … CA – 3’) ---🡪 **Go to 5 -7 (LTR)**
2) The 5’ and 3’ ends of the alignable regions or consensus sequence is: a) a non-coding region, b) the 3’ end display either: Poly-A tails, tandem repeat (microsatellite), or A-rich region, and c) might show truncation at the 5’ end ---🡪 **Go to 8-9 (LINEs Cryptons)**
3) The 5’ and 3’ ends of the alignable regions or consensus sequence has a diagnostic feature in non-coding region ---🡪 **Go to 10-11 (SINEs + Helitrons)**
4) The 5’ and 3’ ends of the alignable regions or consensus sequence has terminal inverted repeats (TIRs) (e.g., 5’ – TAATCTGT … ACAGATTA – 3’) ---🡪 **Go to 12-19 (DNA transposons + DIRs)**
5) TSD is between 4-6 bp:

5.a: Consensus sequence homology shows these protein coding domains =

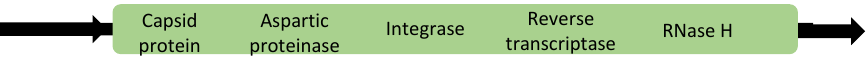
= *Copia* (LTR)

5.b: Consensus sequence homology shows these protein coding domains =

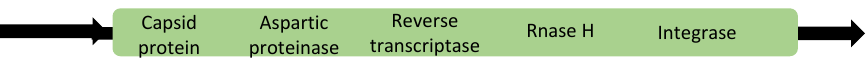
= *Gypsy* (LTR)

5.c: Consensus sequence homology shows these protein coding domains =

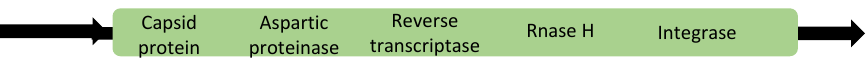
= *Bel-Pao* (LTR)

5.d: Consensus sequence homology shows these protein coding domains =

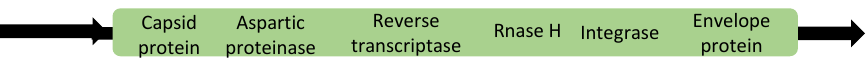
= *Retrovirus* (LTR)

5.e: Consensus sequence homology = shows these protein coding domains

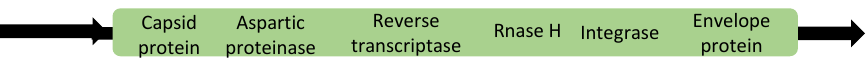
= *EVR* (LTR)

6) Does not have TSD:

8.a: Consensus sequence homology shows these protein coding domains =

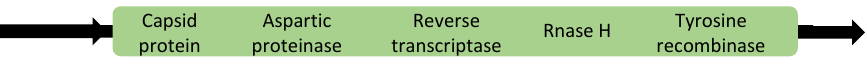
= *Ngaro* (DIRs)

8.b: Consensus sequence homology shows these protein coding domains =

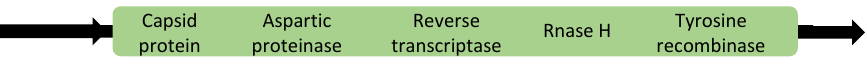
=*Viper* (DIRs)

7) Variable number of bps for TSD and the Consensus sequence homology =

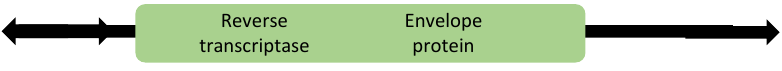

= *Penelope* (PLE)

8) Variable number of bps for TSD:

8.a: Consensus sequence homology shows these protein coding domains =

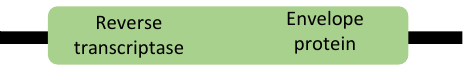

= *R2* (LINEs)

8.b: Consensus sequence homology shows these protein coding domains =

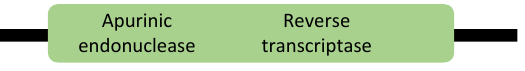

= *RTE* (LINEs)

8.c: Consensus sequence homology shows these protein coding domains =

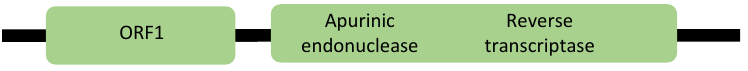

= *Jockey* (LINEs)

8.d: Consensus sequence homology shows these protein coding domains =

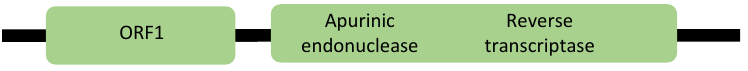

= *L1* (LINEs)

8.e: Consensus sequence homology shows these protein coding domains =

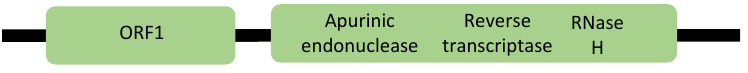

= *I* (LINEs)

9) Does not have TSD and the Consensus sequence homology =

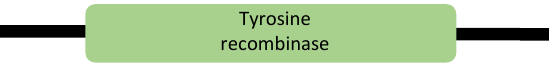
= *Crypton* (Crypton- **DNA transposons**)

10) Variable number of bps for TSD:

10.a: Consensus sequence homology =

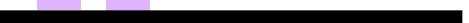

= *tRNA* (SINEs)

10.b: Consensus sequence homology =

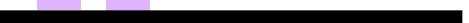

= *7SL* (SINEs)

10.c: Consensus sequence homology =

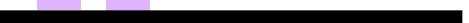

= *5S* (SINEs)

11) Does not have TSD and the Consensus sequence homology =

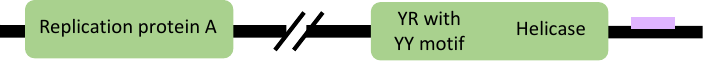

= *Helitron* (Helitron *-* **Rolling circle**)

12) Does not have TSD and the Consensus sequence homology =

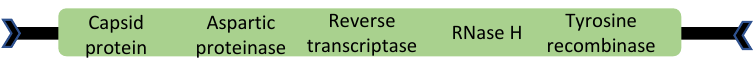
= *DIRs* (DIRs)

13) TSD is “TA” =

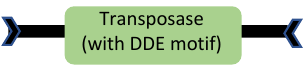

= *Tc1-Mariner*

14) TSD is “TTAA” =

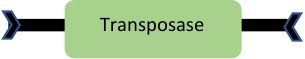

= *PiggyBac*

15) TSD is between 2-3 bp:

13.a: Consensus sequence homology shows these protein coding domains =

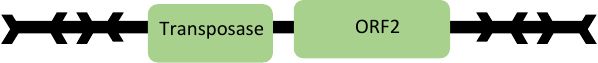

= *CACTA*

13.b: Consensus sequence homology shows these protein coding domains =

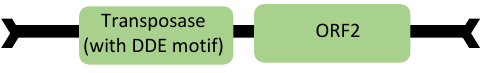

= *PIF – Harbinger*

16) TSD is 5 bp =

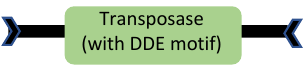

= *Transib*

17) TSD is 6 bp =

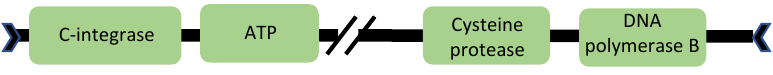

= *Maverick*

18) TSD is between 8-9 bp:

18.a: Consensus sequence homology shows these protein coding domains =

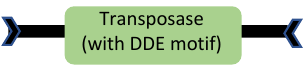

= *hAT*

18.b: Consensus sequence homology shows these protein coding domains =

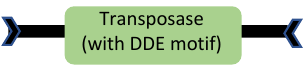

= *Merlin*

18.c: Consensus sequence homology shows these protein coding domains =

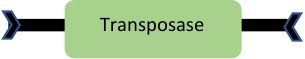

= *P element*

19) TSD is between 9-11 bp =

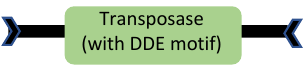

= *Mutator*

**Extra points to validate the identified TE:**

- **DIRs** = contain recombinase gene instead of an INT, therefore do not form TSDs. Their termini are unusual, they resemble either a split direct repeats (SDR) or inverted repeats.
- **PLE** = Encodes and RT that is more closely related to telomere than to the RT from LTR retrotransposons or LINEs and endonuclease that is related both to intron-encoded endonuclease.
- **LINEs** = autonomous LINEs encode an RT and nuclease in their Pol ORF for transposition. Only “I” contain RNase H. The gag-like ORF is sometimes found 5’ to Pol. TSDs are hard to find due to truncation at the 5’ end. At the 3’ end display either; 1) Poly A tail, 2) tandem repeats, and 3) an A-rich region.
- **SINEs** = are mostly Non-autonomous SINEs. Are small (~ 80-500 bp). Generate TSDs that are (5-15 bp). The head which harbors the Pol III promoter defines SINE. The 3’ region can either be; 1) A or AT rich, 2) harbors 3-5 bp tandem repeats, or 3) contain poly (T) tail. Pol III termination signal. The best-known SINE is the human “Alu” element.
- **hAT =** Have TSD of 8 bp, relatively short TIRs of about 5-27 bp, and overall lengths of less that 4 kb.
- **Mutator =** TIRs can extend to several hundred bps, and it has TSDs that is between 9-11 bps.
- **Merlin =** TIRs range from a few dozen to several hundred bp and are flanked by 8-9 bps of TSDs. TSD = 8/9 bp, Length in (kb) = 1.4-3.5, TIRs (bp) = 21-462.
- **Transib =** Contain DDE motif. Related to RAG I protein which in involved in V(D)J recombination. TSD = 5 bp, Length in (kb) = 3-4, TIRs (bp) = 9-60.
- **PiggyBac =** Favors insertion adjacent to “TTAA”. TSD = “TTAA”, Length in (kb) = 2.3-6.3, TIRs (bp) = 12-19.
- **PIF-Harbinger** = Has a target site preference for “TAA”. The TE contains 2 ORFs; one ORF encodes DNA binding protein, the other encodes a DDE transposase. TSD = TWA, Length in (kb) = 2.3-5.5, TIRs (bp) = 15-270.
- **CACTA =** Contain both a transposase and a 2^nd^ ORF of unclear function. In plant, the short TIRs terminate in highly conserved CACTA (sometimes CACTG motifs) and flank 3 bp TSDs. While in animal and fungi, CCC replaces the CACTA motif and 2 bp TSDs are generated. TIRs often flank complex arrays of subterminal repeats. TSD = 2/3 bp, Length in (kb) = 4.5-15, TIRs (bp) = 10-54.
- **Crypton =** Cryptons contain tyrosine recombinase but lack an RT domain; lack TIRs but generate TSDs.
- **Helitron =** Do not generate TSDs and do not have TIRs but rather short terminal motifs. Ends are defined only by TC or CTRR motifs (R = purine) before the 3’ end. TSD = none, Length in (kb) = 5.5-17, TIRs (bp) = none.
- **Maverick (also known as Polintons)** = Are very large transposons reaching 10-20 kb and are bordered by long TIRs and has coding capacity for multiple proteins, encode up to 11 proteins, most of which are related to double-stranded DNA viruses including a B type DNA polymerase. TSD = 5/6 bp, Length in (kb) = 15-25, TIRs (bp) = 150-700.
- **Penelope** = has a pseudo LTRs and an amino acid motif (GIY – YIG).
- **Tc1/*mariner*** = TSD = TA, Length in (kb) = 1.2-5, TIRs (bp) = 17-1100.
- ***P element* =** TSD = 7/8 bp, Length in (kb) = 3-11, TIRs (bp) = 13-150.
